# Changes in interlimb coordination induced by within-stride changes in treadmill speed

**DOI:** 10.1101/2024.11.27.625740

**Authors:** Brooke L. Hall, Ryan T. Roemmich, Caitlin L. Banks

## Abstract

Human walking involves tightly coordinated movements of the right and left legs. We recently developed and tested a ‘dynamic treadmill walking’ paradigm that changes the treadmill speed within a single step to provide asymmetric training for persons with gait dysfunction. We previously demonstrated that this approach could induce changes in human gait symmetry; however, if this approach is to be used in rehabilitation, we also need to understand how movements of the legs are coordinated to produce these asymmetric gait changes. The goal of this study was to examine the temporal (phase shift) and spatial (center of oscillation difference) aspects of interlimb coordination during dynamic treadmill walking in ten young adults without gait impairment. We found that dynamic treadmill walking drove significant changes in phase shift and center of oscillation difference that were dependent on the timing of the treadmill speed change within the gait cycle. For example, slowing the treadmill during the stance phase extended the double limb support period, and these changes were strongly correlated with a phase shift between the two legs. Accelerating the treadmill late in stance led to extensions in the trailing limb angle that were strongly correlated with changes in the center of oscillation difference. Overall, dynamic treadmill walking can be configured to drive changes in many spatiotemporal, kinematic, and interlimb coordination parameters, creating a variety of options for restoring gait symmetry and targeting aspects of spatial and temporal interlimb coordination in clinical populations with heterogenous patterns of gait asymmetry.

**NEW & NOTEWORTHY:** Dynamic treadmill walking has been previously shown to drive asymmetric gait changes, but little is known about its effect on interlimb coordination. Here, we find that this approach induces significant changes in interlimb coordination (relative to normal walking) in both spatial and temporal domains. Altogether, dynamic treadmill walking offers a customizable gait rehabilitation strategy for asymmetric gait training that can induce changes in interlimb coordination using only a single-belt treadmill.

## INTRODUCTION

Many clinical populations – e.g., persons with stroke^1^, Parkinson’s disease^2^, or cerebral palsy^3^ – walk with gait patterns characterized by disrupted interlimb coordination (i.e., poorly coordinated movements of the left and right legs). Unimpaired gait is characterized by oscillating, anti-phase movements between the two legs: one leg generally moves forward as the other extends backward and vice versa in a bilaterally coordinated manner^4,5^. Damage to the nervous system can affect these well-coordinated movements between the legs such that the oscillations deviate from their typical anti-phase relationship and become asymmetric^4,6^. Restoration of anti-phase interlimb coordination then becomes important because deviations from symmetric, coordinated gait can lead to increased fall risk^7,8^ and elevated energy expenditure^9–11^ that can decrease mobility and community engagement^12^.

Interlimb coordination can be quantified in several ways. Locomotion literature often uses a combination of temporal (e.g., limb phasing) and spatial (e.g., difference in the axes around which each limb oscillates, or “center of oscillation difference”) metrics to measure and understand how the legs are controlled in synchrony^5,13–15^. In the temporal domain, “phasing” can be used to capture the degree to which the leg oscillations are in anti-phase^16^. In the spatial domain, “center of oscillation” describes the axis around which each leg rotates in the sagittal plane as a person walks (i.e., a positive or negative center of oscillation would indicate that the leg rotates around a flexed or extended axis, respectively)^17^. The center of oscillation difference then captures the difference in the axes of rotation of the right and left legs. A difference value near zero is expected for symmetric walking, indicating that both legs rotate about a similar axis. However, both phasing and center of oscillation are often altered in persons with asymmetric gait patterns or when walking under asymmetrically perturbed conditions^16^.

The ability to use gait perturbations to drive changes in interlimb coordination offers an opportunity to leverage these perturbations to improve interlimb coordination in persons with gait impairment. Our laboratory recently developed a ‘dynamic treadmill walking’ approach that can drive changes in gait symmetry by changing the speed of a conventional treadmill within a single stride^18^. Our previous study in young adults without gait impairment revealed that the treadmill speed changes could be delivered at different timepoints within the gait cycle (via pacing of participant heel-strikes using a metronome) to induce customizable asymmetries in several clinically relevant gait parameters^18^. For example, slowing the treadmill immediately following right heel-strike induced asymmetries in step time, peak braking, and propulsion impulse whereas slowing the treadmill late in stance induced asymmetries in step time, leading limb angle, and peak braking force^18^.

It is likely that different patterns of changes in interlimb coordination underlie these various spatiotemporal, kinematic, and kinetic gait asymmetries induced by dynamic treadmill walking. However, given that only one limb is paced by a metronome and the other is unconstrained during this task, it is not yet clear how the coordination patterns between the limbs change with treadmill belt speed changes at different timings throughout the gait cycle. We anticipate that a better understanding of how dynamic treadmill walking affects interlimb coordination could be useful for providing customizable rehabilitation approaches aimed at restoring proper interlimb coordination in persons with heterogeneous asymmetric gait deficits.

The goal of this study was to investigate interlimb coordination across a range of dynamic treadmill walking conditions in young adults without gait impairment. We hypothesized that interlimb coordination would be changed during dynamic treadmill walking (relative to normal walking) as demonstrated by non-zero center of oscillation differences and deviations from anti-phase coupling. More specifically, we hypothesized that these changes in interlimb coordination would differ depending on the timing of the speed change within a gait cycle. In the trials where the leg paced by the metronome (the right leg) stepped onto a fast or accelerating treadmill, we hypothesized that leading limb angles on the right leg would increase and stance times would decrease, resulting in the right leg showing a center of oscillation about a more flexed axis when compared to that of the left leg (and the opposite to be true when the right leg stepped onto a slow or decelerating treadmill). Similarly, we expected that the legs would be more in-phase when participants stepped onto a fast or accelerating treadmill with their paced (right) leg with the right leg leading the left in time (and the right leg to lag the left in time when stepping onto a slow or decelerating treadmill).

## METHODS

Ten young adults without gait impairment participated in this study (2 M, 8 F, age: 25.1±3.0 years). These participants composed one group from a larger, unpublished experiment (16 M, 24 F, age: 25.6±3.5 years, randomly assigned to four groups). This group walked under relatively slow dynamic treadmill conditions (as compared to our previous work^18^) for the purpose of understanding potential translational implications to neurologic populations. All participants reported no current neurological, musculoskeletal, or cardiovascular conditions and provided written, informed consent in accordance with the Declaration of Helsinki. The Johns Hopkins School of Medicine Institutional Review Board approved all study procedures, and participants received monetary compensation for their participation (IRB00394009).

### Data collection

Participants walked on an instrumented split-belt treadmill (Motek Medical BV, Houten, The Netherlands), but we emphasize that the treadmill belt speeds were tied for all trials (i.e., participants did not walk under split-belt treadmill conditions with the belts moving at different speeds simultaneously; the split-belt treadmill allowed us to collect ground reaction force data from each individual leg). We collected kinematic data using an eight-camera motion capture system (Vicon Motion Systems, Centennial, CO; 100 Hz; RRID: SCR_015001) with 30 passive retroreflective markers on the seventh cervical vertebrae, jugular notch, tenth thoracic vertebrae, xiphoid process, bilaterally over the iliac crest, anterior superior iliac spine, posterior superior iliac spine, trochanter, thigh, lateral femoral epicondyle, medial femoral epicondyle, shank, lateral malleolus, medial malleolus, calcaneus, second metatarsal head, and fifth metatarsal head. We collected triaxial ground reaction forces using force plates embedded independently under each treadmill belt (1000 Hz). We executed dynamic treadmill walking on our treadmill with a custom D-Flow script (Motek Medical BV, Houten, The Netherlands). The dynamic treadmill controller changed between a slow (0.5 m/s) and fast (1.0 m/s) speed within a single stride at an acceleration rate of 6.0 m/s^2^. For each dynamic treadmill trial, 50% of the gait cycle was spent at the fast speed and 50% at the slow speed.

### Experimental protocol

The experimental protocol is outlined in Figure 1. All participants completed three two-minute baseline trials at the slow (0.5 m/s), intermediate (0.75 m/s), and fast speeds (1.0 m/s). Four five-minute dynamic treadmill walking trials (Slow, Fast, Accelerate, and Decelerate) followed the baseline trials in a randomized order. Our experimental design included a similar set of conditions that we used in our previous work^18^, but here we tested at slower treadmill speeds (previous study: slow=0.75 m/s, intermediate=1.125 m/s, fast=1.5 m/s)^18^. We instructed participants to synchronize each right heel-strike with a metronome tone that was aligned with the treadmill speed changes. We determined the timing of the metronome tone by measuring each participant’s average stride time across the intermediate baseline trial, as calculated by dividing the trial duration (120 seconds) by the total number of strides completed in the trial.

**Figure 1.**
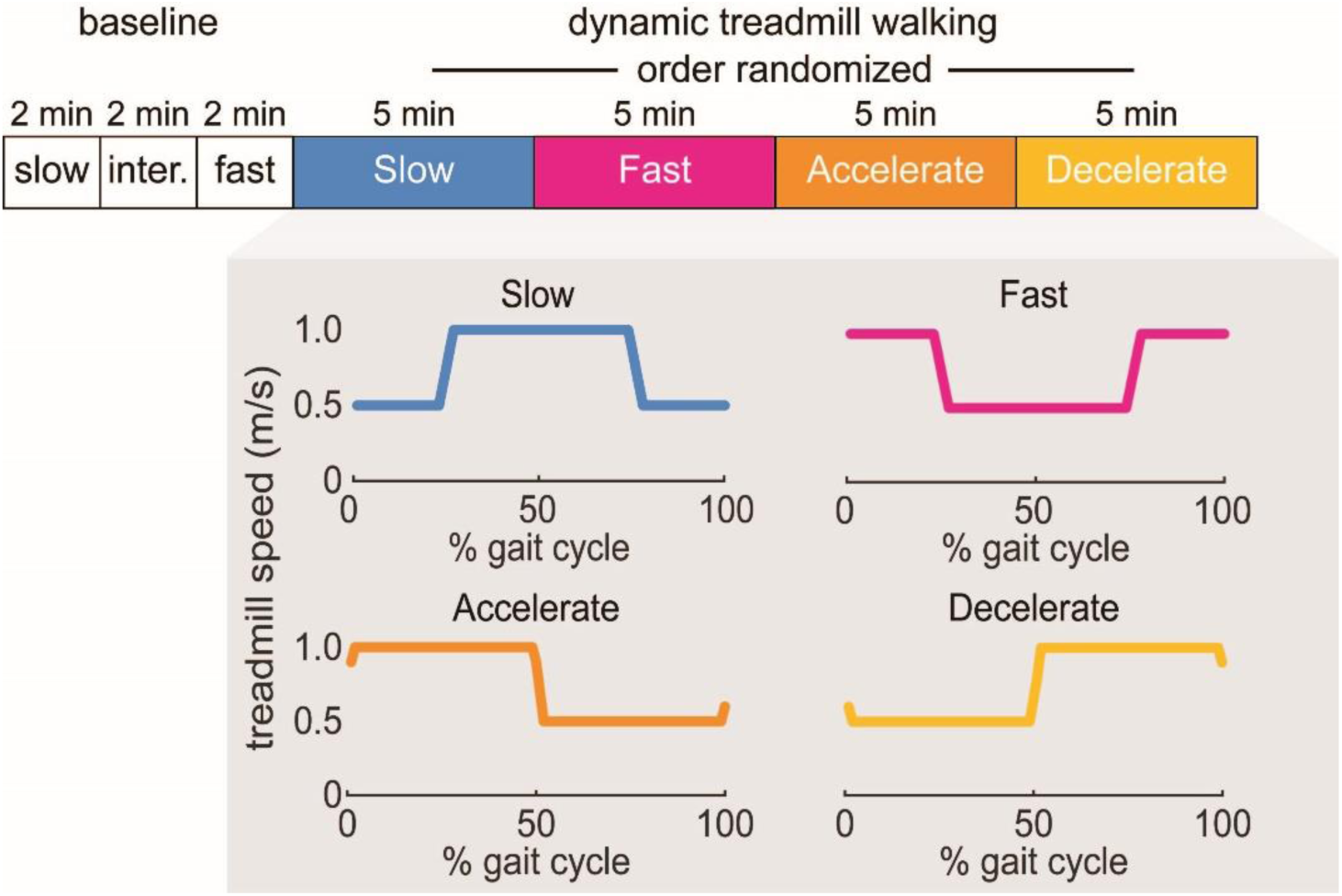
Experimental paradigm: All participants completed three baseline trials at slow (0.5 m/s), intermediate (*inter*, 0.75 m/s), and fast (1.0 m/s) treadmill speeds for two minutes each. Participants then completed the four five-minute, metronome-paced dynamic treadmill walking trials (*Slow, Fast, Accelerate, Decelerate)* in a randomized order. The shaded, gray box depicts the target speed profiles of each trial over a gait cycle. Each trial is named according to treadmill speed at heel-strike (i.e., Slow is at slow treadmill speed at right heel-strike, Fast is at the fast speed, Accelerate is accelerating between slow and fast, and Decelerate is decelerating between fast and slow).

For each dynamic treadmill trial, the treadmill speed changed at different points within the gait cycle (Figure 1). The metronome sounded to pace the participant’s right heel-strike to guide the desired timing pattern set by each dynamic treadmill trial:

- <u>Slow</u>: the right heel-strike was paced to occur with the treadmill at the slow speed (0.5 m/s). This resulted in the treadmill moving slow over the first and last ∼25% of the gait cycle and fast over the middle ∼50% of the gait cycle.
- <u>Fast</u>: the right heel-strike was paced to occur with the treadmill at the fast speed (1.0 m/s). This resulted in the inverse of Slow: the treadmill moved fast over the first and last ∼25% of the gait cycle and slow over the middle ∼50% of the gait cycle.
- <u>Accelerate</u>: the right heel-strike was paced to occur when the treadmill was moving at approximately the intermediate speed (0.75 m/s) while accelerating to the fast speed. This resulted in the treadmill moving at the fast speed during the first ∼50% of the gait cycle and then decelerating to the slow speed for the last ∼50% of the gait cycle
- <u>Decelerate</u>: the right heel-strike was paced to occur when the treadmill moved at approximately the intermediate speed (0.75 m/s) while decelerating to the slow speed. This resulted in the inverse of Accelerate: the treadmill moved slow over the first ∼50% of the gait cycle and fast over the last ∼50% of the gait cycle.

### Data analysis

We used custom MATLAB software for data processing (R2023b, The Mathworks, Natick, MA; RRID: SCR_001622). We filtered the marker and ground reaction force data using fourth-order low-pass Butterworth filters with cutoff frequencies of 6 Hz and 28 Hz, respectively. We determined heel-strikes as the local maxima and minima where the change in limb angle (calculated as the sagittal vectors between the iliac crest and second metatarsal head markers) shifted from positive to negative and toe-offs as the opposite^18^. We calculated step length, step time, and leading and trailing limb angles as in our previous work^18^ as well as stance time (time between heel-strike and toe-off of the ipsilateral limb), swing time (time between toe-off and heel-strike of the ipsilateral limb), and double support time (time between heel-strike of the contralateral limb and toe-off of the ipsilateral limb).

We also recorded the treadmill belt speeds within each trial to assess how well participants synchronized their right heel-strikes with the metronome tone to assess adherence to the desired heel-strike/speed change pairings of the different dynamic treadmill trials. We calculated cross-correlation coefficients (at time lag zero) between the prescribed and actual treadmill speed time series for each gait cycle within each participant to understand how well participants adhered to the prescribed conditions. We also calculated a general metric of how accurately participants stepped to the metronome: for each gait cycle, we considered a participant’s step successful if their right heel-strike occurred within ±200 ms of the metronome tone initiation.

We calculated phase shift and center of oscillation difference as temporal and spatial measures of interlimb coordination, respectively. As a first step to calculating phase shift, we calculated limb phasing as the stride-by-stride maximum cross-correlation coefficient between right and left limb angle trajectories. Next, we subtracted limb phasing during dynamic treadmill walking from limb phasing during intermediate baseline walking to calculate phase shift (i.e., a phase shift of zero would indicate that a participant had identical phasing values to the baseline trial). We calculated center of oscillation as the limb angle at midstance (or at the timepoint midway between heel-strike and toe-off) for each leg. We then calculated the center of oscillation difference as the difference between centers of oscillation measured for the right and left centers of oscillation.

### Statistical analysis

We used R version 4.3.3 (R Core Team, 2023: RRID: SCR_001905) and RStudio^19^ version 2023.12.1 (RRID: SCR_000432) for data analysis and visualization, including the readxl (RRID: SCR_018083), dplyr (RRID: SCR_016708), rstatix (RRID: SCR_021240), and psych (RRID: SCR_021744) packages^19–24^. Given that we paced all participants to their stride time during the intermediate baseline trial, we first used one-way repeated measures ANOVA to confirm that stride time was consistent across the four dynamic treadmill trials. Next, we used one-way repeated measures ANOVA to assess whether order effects were present for our primary interlimb coordination outcome variables (phase shift and center of oscillation difference) across the different trial orders. To test how well participants matched the prescribed belt speeds across the four dynamic treadmill trials, we then performed one-way repeated measures ANOVAs to compare cross-correlation coefficients (averaged across all gait cycles for each participant) of the prescribed versus actual treadmill speed time series across the four dynamic treadmill trials.

To test our primary study hypotheses, we performed a 4 (Trial) x 2 (Leg) repeated measures ANOVA to compare Trial and Leg differences for spatiotemporal and kinematic gait parameters (i.e., stance time, swing time, double support time, leading limb angle, and trailing limb angle). We then investigated interlimb coordination across the dynamic treadmill trials using one-way repeated measures ANOVA to compare phase shift among four dynamic treadmill trials and performed a similar analysis on center of oscillation difference. We analyzed individual leg centers of oscillation using 4 (Trial) x 2 (Leg) repeated measures ANOVA to understand whether any center of oscillation differences were driven by changes in center of oscillation in the right versus left leg. All repeated measures ANOVAs were tested for sphericity using Mauchly’s Test, and Greenhouse-Geisser corrected values are provided when the sphericity test failed (p<0.05)^25^. Finally, we performed Pearson’s correlations to assess the relationships between the interlimb coordination measures and kinematic and spatiotemporal gait parameters. For all intralimb and interlimb measures, we performed statistical analyses on the mean of the last 30 strides for each trial. We set α<0.05 for all analyses and used Bonferroni post-hoc corrections for pairwise comparisons where appropriate.

## RESULTS

Our preliminary checks verified that 1) participants walked with similar stride times across all four dynamic treadmill walking trials and 2) there was no effect of the dynamic treadmill trial order on the outcomes. We verified that participants indeed walked with similar stride times across all four dynamic treadmill walking conditions using one-way repeated measures ANOVA, where there was no significant main effect of Trial (F(3,67)=2.73, p=0.05). Similarly, we did not observe a significant main effect of Block (or order) on phase shift (F(3,27)=0.92, p=0.45) or center of oscillation difference (F(3,27)=0.41, p=0.75).

### Participants stepped in synchrony with the metronome during dynamic treadmill walking

Participants synchronized their right heel-strikes to a metronome tone with strong cross-correlation coefficients (at time lag zero) between the prescribed and actual treadmill speeds across each of the trials (mean±standard error of the mean): Slow: r=0.97±0.01, Fast: r=0.98±0.01, Accelerate: r=0.98±0.01, Decelerate: r=0.97±0.01(treadmill speed profiles shown in Figure 2A). One-way repeated measures ANOVA revealed no significant main effect of Trial (F(3,26)=1.59, p=0.21) on the cross-correlation coefficients, indicating that there were no significant differences in how successfully participants stepped to the metronome tone across the four dynamic treadmill trials. Moreover, participants were generally successful with synchronizing their right heel-strikes to the metronome tone: group mean success rates ranged from 89% to 94% across the four dynamic treadmill trials (i.e., participants stepped within ±200 ms of the metronome tone on 89% to 94% of strides).

**Figure 2.**
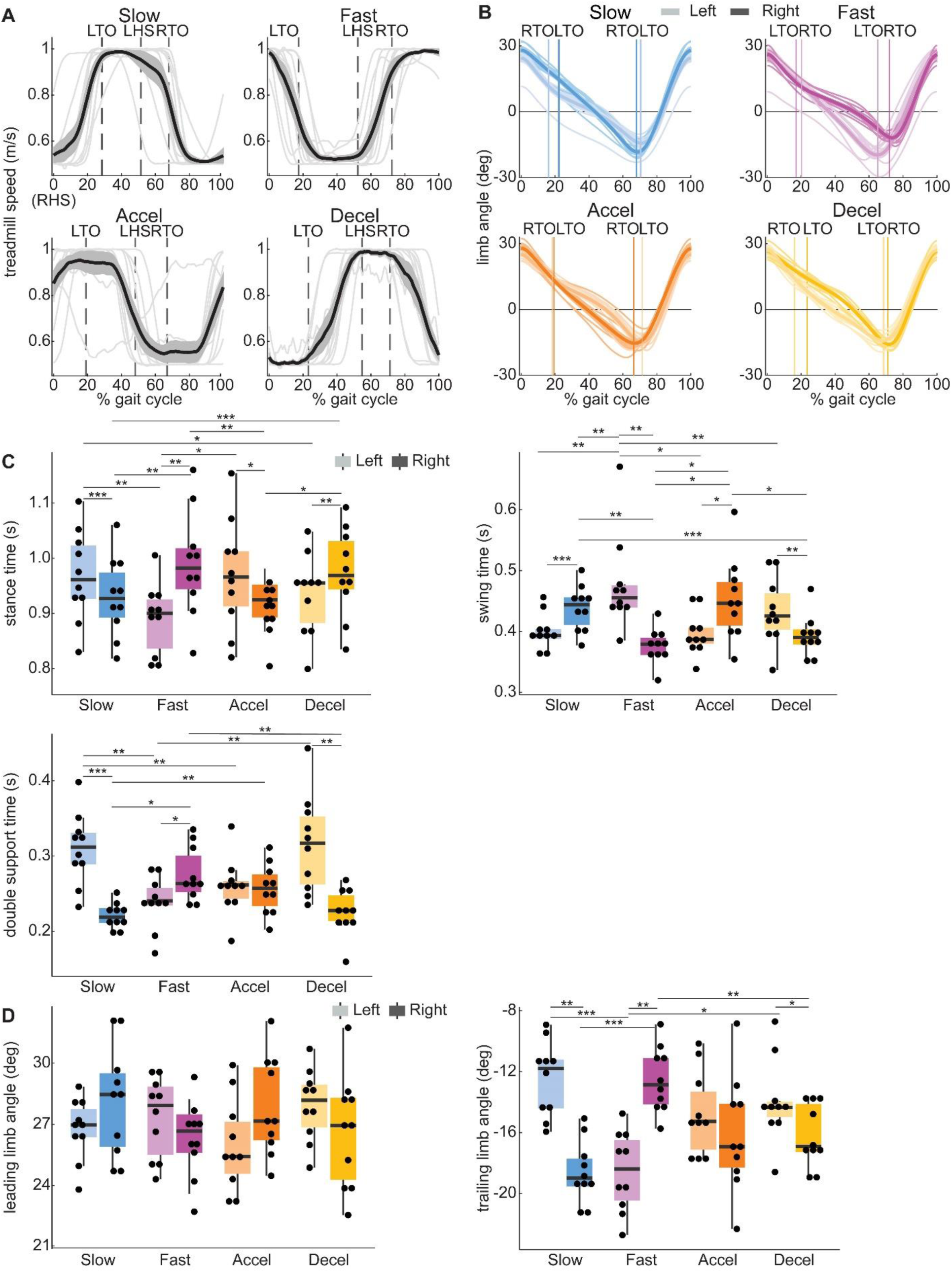
Group speed profiles, limb angles, and spatiotemporal gait parameters: **A)** Group average treadmill speed traces are shown as solid black lines with standard error of the mean represented with gray fill. Individual participant speed profiles are layered behind the group traces in light gray. The group average gait event timings (LTO, LHS, RTO) are represented by vertical dashed lines shown in each profile. *RHS*: right heel-strike; *LTO:* left toe-off; *LHS:* left heel-strike; *RTO:* right toe-off. The four speed conditions include *Slow, Fast, Accelerate* (*Accel*), and *Decelerate* (*Decel*). **B)** Limb angle plots with the right (dark) and left (light) gait cycles anchored to their own heel-strike (i.e., 0% of the gait cycle), respectively. Line conventions and abbreviations follow those in Figure 2A. **C and D)** Box and whisker plots for spatiotemporal (**C**) and kinematic (**D**) parameters. The black scatter points show individual average values over the last 30 strides for each participant. The horizontal lines indicate statistical significance as denoted with asterisks (*: p<0.05, **: p<0.01), ***: p<0.001). Shading conventions follow Figure 2B.

### Dynamic treadmill walking drives asymmetric changes in spatiotemporal and kinematic gait parameters

Next, we aimed to understand the spatiotemporal and kinematic factors that contributed to the different patterns of limb angle timing and amplitude observed across the four dynamic treadmill trials (Figure 2B). For comparison to our previous work, we calculated step length in addition to the step-by-step values of spatiotemporal and kinematic parameters below (Supplemental Figures S1-S4).

#### Stance time

Stance time generally increased when the treadmill was slow during a longer portion of the stance phase (e.g., right leg stance during the Fast and Decelerate conditions). A 4 (Trial) x 2 (Leg) repeated measures ANOVA revealed a significant main effect of Trial (F(3,27)=3.44, p=0.03) on stance time and a significant Trial*Leg interaction (F(3,27)=14.3, p<0.01), but no significant main effect of Leg (F(1,9)=2.09, p=0.18; Figure 2C). Post-hoc analyses showed significant within-trial Leg differences for stance time in Slow (left longer than right, p<0.01), Fast (right longer than left, p<0.01), Accelerate (left longer than right, p=0.04), and Decelerate (right longer than left, p<0.01). We also observed that left stance time in Fast was significantly shorter than left stance time in Accelerate (p=0.03) and Slow (p<0.01). Right stance time in Slow was significantly shorter than in Decelerate (p=0.01) and Fast (p<0.01). Left stance time was significantly longer in Slow than in Decelerate (p=0.046). Right stance time in Accelerate was significantly shorter than in Decelerate (p=0.01) and Fast (p<0.01).

#### Swing time

Because swing time has a strong, inverse correlation with stance time across all trials (r^2^=-0.99, p<0.01), we expected similar findings as stance time but with opposite directionality. Indeed, swing times generally decreased when the treadmill was slow during a large portion of the stance phase and fast during the swing phase. A 4 (Trial) x 2 (Leg) repeated-measures ANOVA revealed no significant main effect of Trial (F(3,27)=2.85, p=0.06) or Leg (F(1,9)=2.07, p=0.18) on swing time but did reveal a significant Trial*Leg interaction (F(3,27)=14.55, p<0.01, Figure 2C). Post-hoc analyses revealed significant within-trial Leg differences in swing time for Slow (right longer than left, p<0.01), Fast (left longer than right, p<0.01), Accelerate (right longer than left, p=0.04), and Decelerate (left longer than right, p<0.01). Furthermore, post-hoc analyses revealed that within-leg swing times were significantly different between Fast and Accelerate, with Fast producing a longer swing time on the left leg than Accelerate (p=0.02) and Accelerate driving a longer swing time on the right leg (p=0.01). Slow and Fast exhibited significantly different swing times on the left and right legs, with Fast producing a longer left swing time than Slow (p=0.01) and Slow producing a longer right swing time than Fast (p<0.01). Decelerate produced a significantly shorter right swing time than Accelerate (p=0.02) and Slow (p<0.01).

#### Double support time

Double support times were generally longer when the treadmill was slow during double support (e.g., double support period from right heel-strike to left toe-off – or left double support time using our naming convention – during Slow or Decelerate). A 4 (Trial) x 2 (Leg) repeated measures ANOVA revealed a significant main effect of Trial (F(3,27)=3.31, p=0.04) and Leg (F(1,9)=8.62, p=0.02) on double support time as well as a significant Trial*Leg interaction (F(3,27)=13.17, p<0.01). Post-hoc analyses revealed significant Leg differences within Slow (left longer than right, p<0.01), Fast (right longer than left, p=0.047), and Decelerate (left longer than right, p<0.01). Left double support time in Fast was significantly shorter than left double support time in Decelerate (p<0.01) and Slow (p<0.01). Left double support time in Slow was significantly longer than that in Accelerate (p<0.01). Slow produced significantly shorter right double support times than Accelerate (p<0.01) and Fast (p=0.01). Right double support time was significantly longer in Fast than in Decelerate (p<0.01).

#### Leading limb angle

Leading limb angles were largely similar across trials. A 4 (Trial) x 2 (Leg) repeated measures ANOVA revealed no significant main effect of Trial (F(3,27) =0.81, p=0.50) or Leg (F(1,9)=1.61, p=0.24) on leading limb angle. We also did not observe a significant Trial*Leg interaction (F(3,27)=2.27, p=0.10).

#### Trailing limb angle

Trailing limb angles were generally larger (i.e., more negative) when the treadmill moved fast during late stance (e.g., right leg in Slow). A 4 (Trial) x 2 (Leg) repeated measures ANOVA revealed no significant main effects of Trial (F(3,27)=1.56, p=0.22) or Leg (F(1,9)=0.19, p=0.67) on trailing limb angle, but we did observe a significant Trial*Leg interaction (F(3,27)=12.66, p<0.01). Post-hoc analyses revealed that Slow, Fast, and Decelerate showed significant differences between the left and right legs, with right trailing limb angle being larger (i.e., more negative) than left trailing limb angle in Slow (p<0.01), the left trailing limb angle being larger than the right trailing limb angle in Fast (p<0.01), and right trailing limb angle being larger than left trailing limb angle in Decelerate (p=0.049). Left trailing limb angle in Fast was significantly larger than in Decelerate (p=0.03) and Slow (p<0.01). Right trailing limb angle in Fast was significantly smaller than in Decelerate (p<0.01) and Slow (p<0.01).

### Dynamic treadmill walking alters spatial and temporal components of interlimb coordination

#### Temporal interlimb coordination: phase shift

After understanding how the different dynamic treadmill trials affected spatiotemporal and kinematic gait parameters, we next studied how the different trials affected interlimb coordination. A phase shift value of zero indicates that the limbs are in anti-phase (similar to baseline walking). Positive phase shift indicates that the right leg oscillations are advanced in time relative to those of the left and vice versa. In Figure 3A, we now show the limb angles with 0% of the gait cycle representing right heel-strike to visualize the coordination between the two limbs.

**Figure 3.**
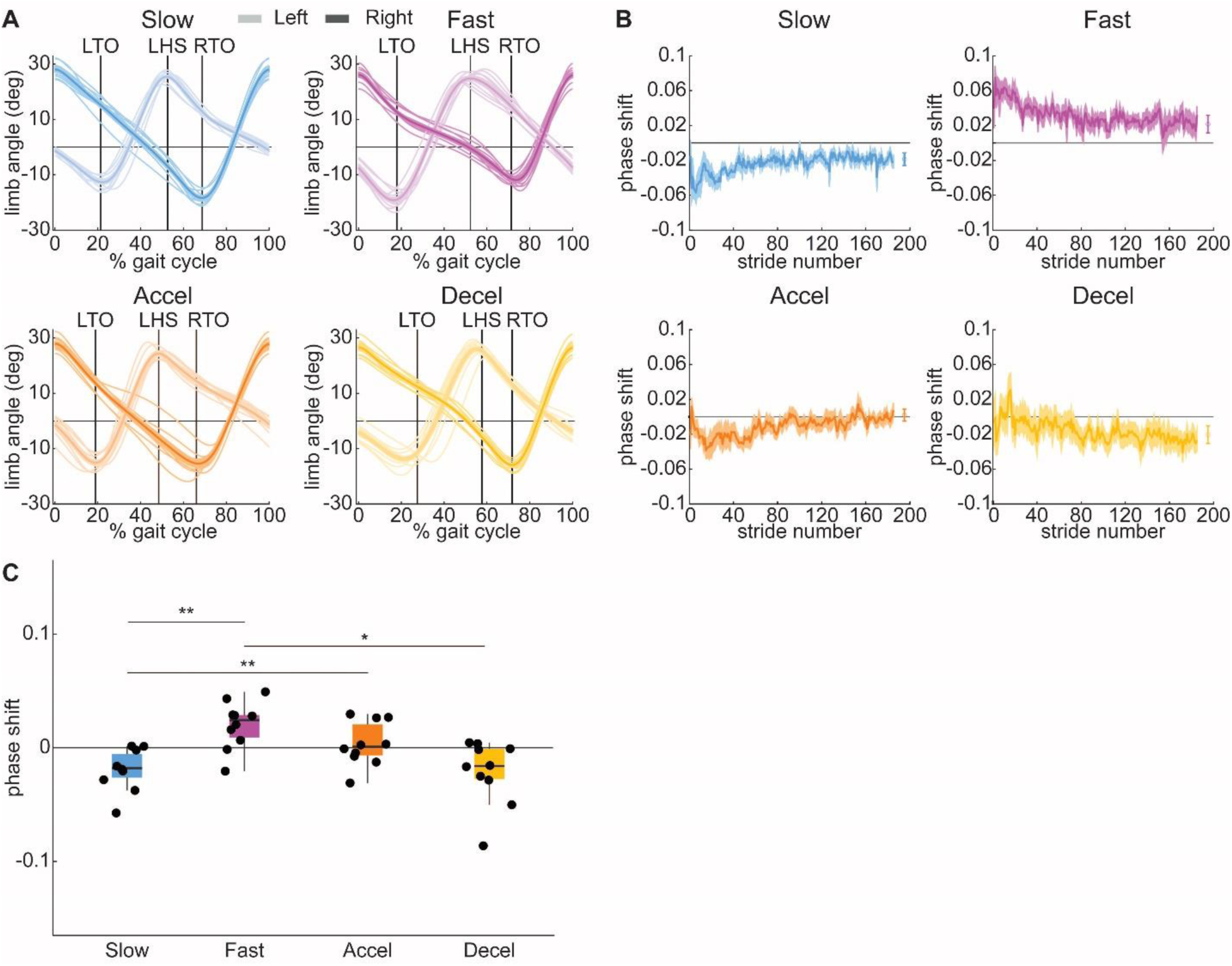
Phase shift: **A)** Limb angle plots anchored to the right heel-strike. Lines and shading conventions follow Figure 2B. Individual participant traces for each leg are displayed by the thin lines. Group averages of each gait event for each gait cycle are represented by vertical solid lines. **B)** Time-series plots of left and right limb angles through the duration of each dynamic treadmill walking condition. Plots depict group mean (thick line) ± standard error of the mean (lighter colored fill). On the far-right side of each plot, group mean ± standard error over the last 30 strides is depicted by a solid dot with error bar. **C)** Box and whisker plots for phase shift follow the same conventions as Figures 2C and 2D. The black scatter points show individual average values over the last 30 strides for each participant. The horizontal lines indicate statistical significance as denoted with asterisks (*: p<0.05, **: p<0.01), ***: p<0.001).

Dynamic treadmill walking conditions showed the most pronounced changes in phase shift when the treadmill speed changes occurred largely during double support periods (e.g., Slow and Fast, see Figures 3B-C). One-way repeated measures ANOVA revealed a significant main effect of Trial (F(2.2,19.76)=9.7, p<0.01) on phase shift (Figure 3C). Post-hoc analyses revealed that phase shift in Fast was significantly more positive (i.e., right leg oscillates advanced in time relative to the left) than in Slow (p<0.01) and Decelerate (p=0.02). Phase shift in Slow was significantly more negative (i.e., left leg oscillates advanced in time relative to the right) than in Accelerate (p<0.01).

#### Spatial interlimb coordination: center of oscillation and center of oscillation difference

A positive center of oscillation value indicates that the right leg is rotating about a flexed axis, whereas a negative value indicates rotation about an extended axis. For center of oscillation difference, a positive value indicates that the right leg rotates about a more flexed axis than the left and vice versa.

Similar to our findings in the temporal domain, dynamic treadmill walking conditions where the treadmill speed changes occurred largely during double support periods showed the most pronounced changes in center of oscillation (Figure 4). A 4 (Trial) x 2 (Leg) repeated measures ANOVA revealed no significant main effects of Trial (F(3,27)=0.85, p=0.45) or Leg (F(1,9)=0.96, p=0.35) on center of oscillation. However, the ANOVA did reveal a significant Trial*Leg interaction (F(3,27)=7.24, p<0.01; Figure 4C). Post-hoc analyses revealed that center of oscillation in Fast was significantly more positive on the right leg than the left (p=0.02). Slow produced a significantly more positive center of oscillation on the left leg than the right (p<0.01). Similarly, Decelerate drove a significantly more positive center of oscillation on the left leg than the right (p=0.03). In Fast, the left leg oscillated about a more extended axis when compared to the left leg in Decelerate (p<0.01). In Slow, the left leg oscillated about a more flexed axis when compared to the left leg in Accelerate (p=0.049). In Fast, the left leg oscillated about a more extended axis than the left leg in Slow (p<0.01). Conversely, the right leg in Fast oscillated about a significantly more flexed axis when compared to the right leg in Slow (p=0.03).

**Figure 4.**
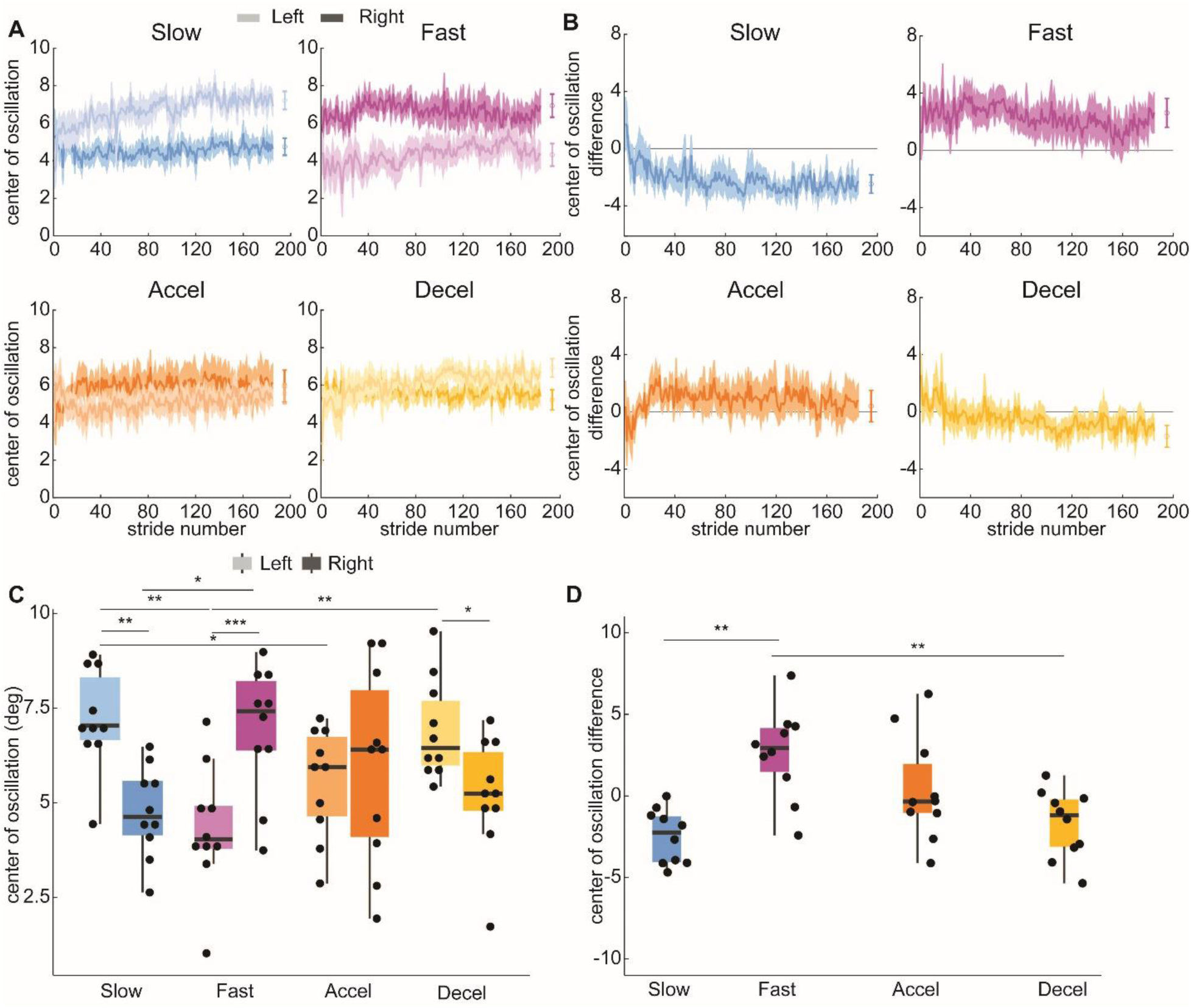
Center of oscillation: **A)** Time-series plots of the spatial shift between the left (lighter color) and right (darker color) limb angles through each dynamic treadmill trial. Plots depict group mean (thick line) ± standard error of the mean (lighter fill). On the far-right side of each plot, group mean ± standard error over the last 30 strides is depicted by a solid dot with error bar. **B)** Time series plots of center of oscillation difference for each walking trial. **C)** Boxplots of center of oscillation from each trial. Boxes display the group median ± interquartile range of phase shift values. Individual participant data are represented as black scatter points. **D)** Boxplots of center of oscillation difference for each trial. Asterisks indicate statistical significance within a repeated measures analysis of variance: *: p<0.05, **: p<0.01, ***: p<0.001.

For center of oscillation difference (Figure 4D), a one-way repeated measures ANOVA revealed a significant main effect of Trial (F(3,27)=7.24, p<0.01). Post-hoc analyses revealed that Fast displayed a significantly larger center of oscillation difference than Decelerate (p<0.01). Additionally, Fast displayed a significantly larger center of oscillation difference than Slow (p<0.01).

### Correlational relationships between interlimb gait parameters and spatiotemporal and kinematic gait parameters

#### Phase shift

We observed several statistically significant correlations between phase shift and spatiotemporal and kinematic gait parameters (Figure 5A). We observed a significant positive correlation between phase shift and stance time asymmetry (r=0.41, p<0.01) and a significant negative correlation between phase shift and swing time asymmetry (r=-0.42, p<0.01). We also observed a significant positive correlation between phase shift and double support time (r=0.93, p<0.01). Furthermore, we observed significant negative correlations between phase shift and leading and trailing limb angles (r=-0.33, p=0.03; r=-0.47, p<0.01).

**Figure 5.**
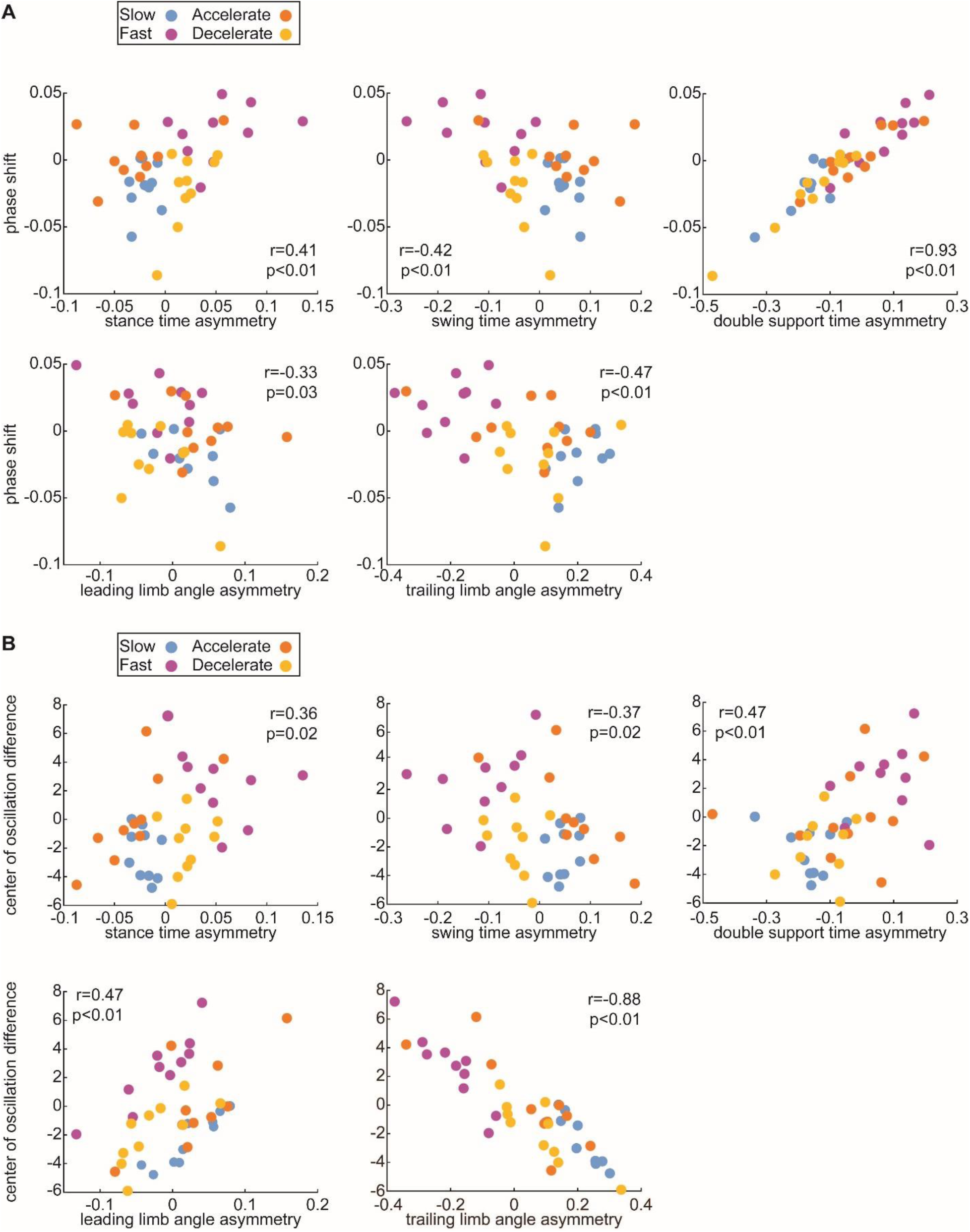
Correlational relationships between spatiotemporal and kinematic asymmetry and intralimb parameters: **A)** Scatter plots of center of oscillation difference versus asymmetry values for stance time (top left), swing time (top center), double support time (top right), leading limb angle (lower left), and trailing limb angle (lower center). **B)** Scatter plots of phase shift versus spatiotemporal (upper) and kinematic (lower) asymmetry values. Pearson correlation coefficients and statistical significance are listed on each plot.

#### Center of oscillation difference

Center of oscillation difference was also significantly associated with spatiotemporal and kinematic gait parameters (Figure 5B). Similar to phasing, we observed significant positive correlations between center of oscillation difference and stance time asymmetry (r=0.36, p=0.02) and double support time asymmetry (r=0.47, p<0.01) as well as a significant negative correlation between center of oscillation difference and swing time asymmetry (r=-0.37, p=0.02). We observed a significant positive correlation between center of oscillation difference and leading limb angle (r=0.47, p<0.01) and a significant negative correlation between center of oscillation difference and trailing limb angle (r=-0.88, p<0.01).

## DISCUSSION

In this study, we showed that dynamic treadmill walking drove significant interlimb coordination changes that were dependent on the timing of the speed change within the gait cycle (i.e., trial type) and were related to a variety of changes in spatiotemporal and kinematic gait parameters. Overall, our hypotheses were supported in Slow and Fast, which were the trials showing the most strongly perturbed interlimb coordination in both the spatial and temporal domains. During Slow and Fast, the legs were more in-phase and showed larger asymmetries in the axes of rotation for each leg when compared to the other dynamic treadmill trials and to normal walking. Here, we will discuss potential implications of the different dynamic treadmill walking trials in driving gait changes in populations with gait asymmetry, and future implications for the potential clinical translation of dynamic treadmill walking.

### Spatiotemporal and kinematic changes depended on the timing of treadmill speed changes

Dynamic treadmill walking drove significant changes in spatiotemporal and kinematic gait parameters. Despite the changing nature of the treadmill speed during dynamic treadmill walking, the spatiotemporal and kinematic changes were in accordance with those that may be expected based on the speeds of the treadmill isolated at specific points in the gait cycle. For example, the treadmill moved at the slow speed for a majority of the stance phase of the right leg during Fast; accordingly, the right stance time was significantly longer than the left stance time during this trial. Similarly, trailing limb angle was influenced by treadmill speed during late stance. A fast treadmill speed during late stance drove a larger (more negative) trailing limb angle. Surprisingly, leading limb angle did not follow our expectations, as we expected the leg that stepped onto a fast treadmill to show a larger leading limb angle than the other leg (i.e., we expected larger leading limb angles in the right leg during Fast as compared to Slow^18^).

However, there were no significant differences between the legs within any or between the trials. Our leading limb angle findings do not agree with our previous work^18^; we hypothesize that this may be due to the slower belt speeds being less likely to alter foot placement than the speeds used previously^18^.

### Changes in interlimb coordination were related to changes in spatiotemporal and kinematic gait parameters

Dynamic treadmill walking induced significant temporal and spatial interlimb coordination changes as quantified by phase shift and center of oscillation difference, respectively. When the treadmill was fast at heel-strike but then decelerated during double support (i.e., Fast), the leading leg rotated about a more flexed axis and progressed through an extended stance phase. This also drove the leading leg to lag in phase relative to the contralateral leg. As expected, opposite interlimb coordination changes were driven between dynamic treadmill walking trials with opposite behavior (e.g., Slow and Fast). Overall, the legs were more in-phase and participants walked about a more forward axis of oscillation than in normal walking.

We also observed that these changes in interlimb coordination were strongly associated with a variety of spatiotemporal and kinematic gait parameters. For instance, we found that double support time asymmetry was strongly correlated with phase shift, indicating that double support periods of the gait cycle were sensitive to treadmill speed change and related to similar shifts in phasing. In the spatial domain, we observed a strong correlation between center of oscillation difference and trailing limb angle asymmetry. Unsurprisingly, when the treadmill accelerated or moved at the faster speed during late stance, the treadmill extended the leg in stance further behind the body than the other, ultimately contributing to the right leg oscillating about a more extended axis of rotation.

### Comparison between dynamic treadmill walking and split-belt walking

Dynamic treadmill walking could be intuited as a split-belt walking-like environment (in split-belt walking, there are separate treadmill belts under each foot that drive each leg to walk at a different speed simultaneously^5^) with a conventional, single-belt treadmill. However, these two approaches to asymmetric gait training have clear differences. During split-belt walking, the belt speeds experienced by the left and right legs are independent; in dynamic treadmill walking, each leg experiences the same belt speed during double support periods. Therefore, in dynamic treadmill walking, the bilateral mismatch in the belt speed predictions that occurs during split-belt treadmill walking is absent.

The interlimb coordination demands of split-belt treadmill walking are also significantly different than those observed here in dynamic treadmill walking. During split-belt walking, the slow leg lags the fast leg in timing, creating a more in-phase gait pattern than normal walking^16^. Additionally, the slow leg oscillates about a more forward axis than the fast leg during split-belt walking^16^. Here, Slow and Fast dynamic treadmill walking conditions drove similar patterns of temporal and spatial changes, but Accelerate and Decelerate produced changes in only one, or neither, domain of interlimb coordination due to the timing of the speed changes. The magnitude of interlimb coordination change created in Slow and Fast was similar to that of gradually introduced split-belt walking but about half the magnitude of abruptly introduced split-belt walking^16^. We hypothesize that this lower magnitude of interlimb coordination change in Slow and Fast – and smaller deviations in the remaining trials – play a role in dynamic treadmill walking not following the characteristics of gait adaptation that are commonly observed in split-belt walking. For example, our previous work showed that there were minimal aftereffects after dynamic treadmill walking^18^, certainly lower than the robust aftereffects observed in split-belt walking^18^. We suspect that these phenomena are driven by the lack of bilateral mismatch in belt speed predictions that occur in split-belt walking. Additionally, the explicit task of pacing the right limb to a metronome provides a different kind of stimulus for gait pattern change than the implicit task of responding to asymmetric belt speed changes alone^26^.

### Clinical translation of dynamic treadmill walking

The ability to selectively target asymmetric gait using dynamic treadmill walking, which can be executed on a conventional treadmill, offers new opportunities for use in traditional rehabilitation settings. To understand how people with a neurologic impairment or injury will respond to dynamic treadmill walking, it is important to first understand the interlimb coordination demands of this pattern. We have shown that this pattern can be leveraged to drive significant interlimb coordination changes in Slow or Fast but does not require the same magnitude of changes as other methods (i.e., split-belt walking^27^). For people with gait impairment, the field is still gaining insights into the necessary ingredients for creating a long-term change in gait symmetry^28^ and perturbing interlimb coordination is one of the potential candidates. Additionally, we cannot assume that people with asymmetric gait patterns, and especially those with neurologic conditions, respond to dynamic treadmill walking in the same way as unimpaired young adults. Therefore, ongoing work is assessing the feasibility of using dynamic treadmill walking to normalize asymmetry patterns after stroke, as well as the potential impact of repeated exposure on changing overground gait asymmetry.

### Limitations

We note several limitations to our study. Young adults with intact neurologic systems tend to walk at faster speeds than studied here (approximately 1.5 m/s^29^); however, in our study, participants walked at 0.5 m/s and 1.0 m/s on the dynamic treadmill. These slower speeds are advantageous from a clinical perspective because clinical populations tend to walk more slowly, though there is potential that these slower speeds may have simultaneously disrupted the normal walking patterns of our participants^30^. Another limitation of the current study is that we only used two measures of interlimb coordination (center of oscillation difference and phase shift).

There are other measures that we could have used to gain a deeper understanding of interlimb coordination changes driven by dynamic treadmill walking (e.g., limb orientation at weight shift^5^). Similarly, we operationally defined interlimb coordination as the coordination between the left and right legs. We therefore did not assess movement of the upper limbs or how they interact with the legs in these analyses. Finally, while our long-term goals of this line of research are focused on clinical applications, we did not study clinical populations directly.

## CONCLUSION

In this study, we investigated the interlimb coordination changes that dynamic treadmill walking may require. We found that dynamic treadmill walking drove significant interlimb coordination changes, both in the spatial and temporal domains. These changes were dependent on the timing of the treadmill speed change. These findings show that dynamic treadmill walking is a tunable tool that can drive changes in spatiotemporal, kinematic, and interlimb coordination parameters. As a potential clinical tool, this finding is encouraging because it offers additional flexibility to remediate gait asymmetry while choosing whether to challenge interlimb coordination and the direction in which it is challenged. Given that we tested interlimb coordination changes driven by this tool in slower-than-average walking for control participants, there is potential for implementation into clinical populations that present with gait asymmetry.

## DATA AVAILABILITY

The data and analysis code that supports the findings of this study are archived on Zenodo: https://doi.org/10.5281/zenodo.15705993.

## ACKNOWLEDGEMENTS

The authors would like to thank the members of the Kennedy Krieger Institute Center for Movement Studies for their valuable contributions and thoughtful discussion to the findings presented here. This article was posted to BioRxiv as a preprint, available at https://doi.org/10.1101/2024.11.27.625740.

## AUTHOR CONTRIBUTIONS

**Conceived and designed research**: BH, RTR

**Performed experiments:** BH

**Analyzed data:** CLB, BH

**Interpreted results of experiments:** CLB, BH, RTR

**Prepared figures:** CLB, BH, RTR

**Drafted manuscript**: BH

**Edited and revised manuscript:** CLB, BH, RTR

**Approved final version of the manuscript:** CLB, BH, RTR

