## Supplementary Figures S1-S4 for "Changes in interlimb coordination induced by within-stride changes in treadmill speed"

### SUPPLEMENTARY MATERIAL

*Relationships between clinically relevant biomechanical and interlimb coordination parameters*

Our previous dynamic treadmill walking study investigated how clinically relevant biomechanical parameters changed within and between the four dynamic treadmill conditions[18]. Our findings are qualitatively similar to this previous work but are included here to assess the relationship between clinically relevant gait parameters and our interlimb coordination parameters at slower treadmill speeds (i.e., 0.5 and 1.0m/s).

A 4 (Trial) x 2 (Leg) repeated measures ANOVA revealed no significant main effect of Trial ($F(1.28,11.48)=1.97$, $p=0.19$) or Leg ($F(1,9)=0.27$, $p=0.61$) on step length. There was a significant Trial*Leg interaction on step length ($F(1.81,16.28)=8.98$, $p=0.03$). Post-hoc analyses revealed that step length was significantly different between the legs in Fast (right larger than left, $p<0.01$) and Slow (left larger than right, $p<0.01$). Step time revealed no significant main effect of Trial ($F(3,27)=1.45$, $p=0.25$), but a significant main effect of Leg ($F(1,9)=22.3$, $p<0.01$). Similarly, step time had a significant Trial*Leg interaction ($F(3,27)=10.3$, $p<0.01$). Post-hoc analyses revealed significant within trial leg differences on step time in Slow (right longer than left, $p<0.01$), Accelerate (left longer than right, $p=0.03$), and Decelerate (right longer than left, $p<0.01$). Left step time in Accelerate was significantly longer than left step time in Decelerate ($p<0.01$) and Slow ($p=0.04$). Right step time in Accelerate was significantly shorter than right step time in Decelerate ($p<0.01$) and Slow ($p=0.02$). Overall, Decelerate produced the greatest magnitude of asymmetry for step time between the left and right legs.

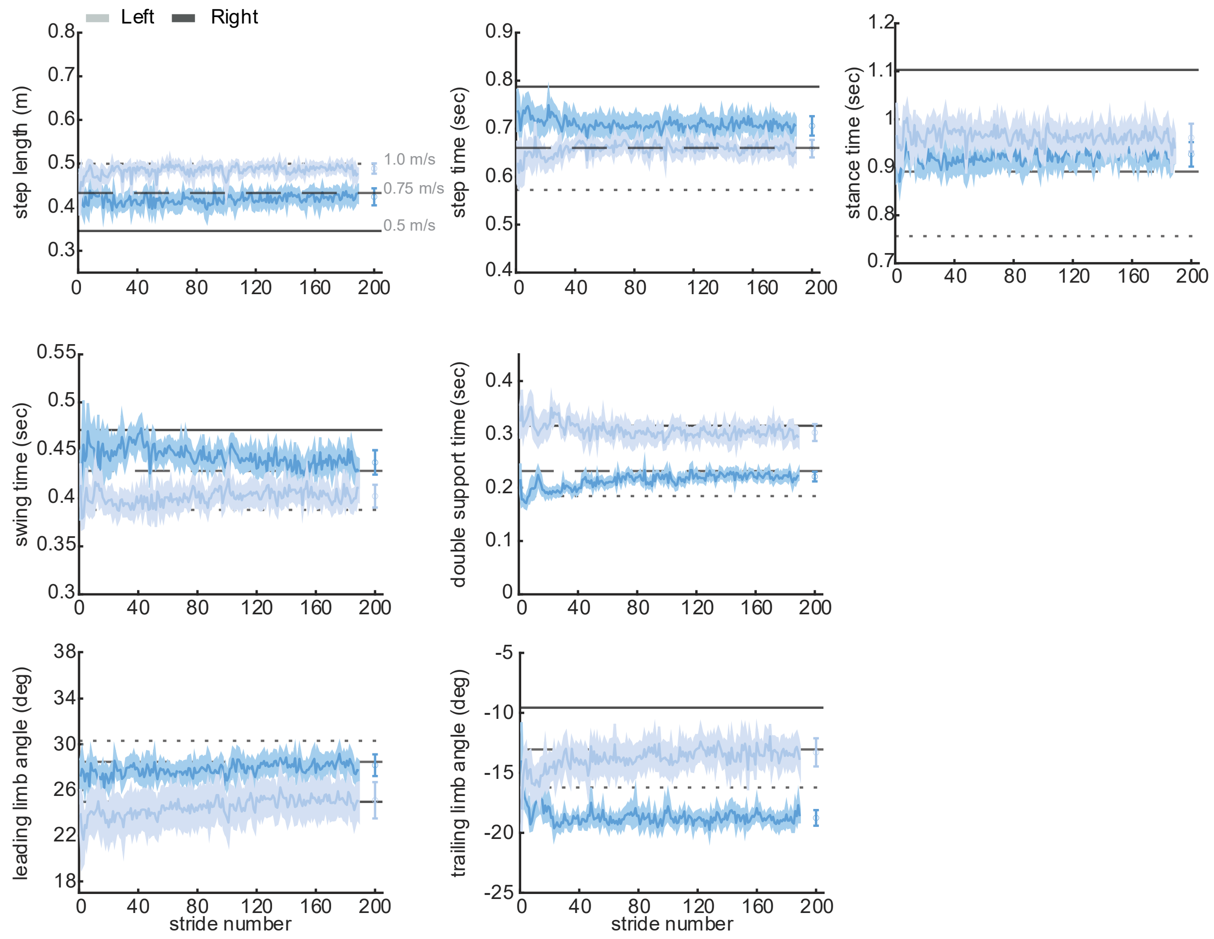


**Figure S1.** Slow spatiotemporal and kinematic traces: Plots depict step-by-step values for step length (top left), step time (top middle), stance time (top right), swing time (second row left), double support time (second row right), leading limb angle (third row right), and trailing limb angle (third row left) within Slow. Right leg is noted by the darker shade and left in the lighter shade. Group mean values are shown by the solid lines, lighter fill surrounding mean lines indicate group standard error of the mean. Scatter points and error bars at the end of each time series trace indicates the group average and standard error over the last 30 strides of the trial. Asymmetry patterns all agree with our previous work[18] but are collected at slower treadmill speeds. Baseline walking values at each speed are shown by horizontal lines (1.0 m/s: dotted; 0.75 m/s: dashed; 0.5 m/s: solid).

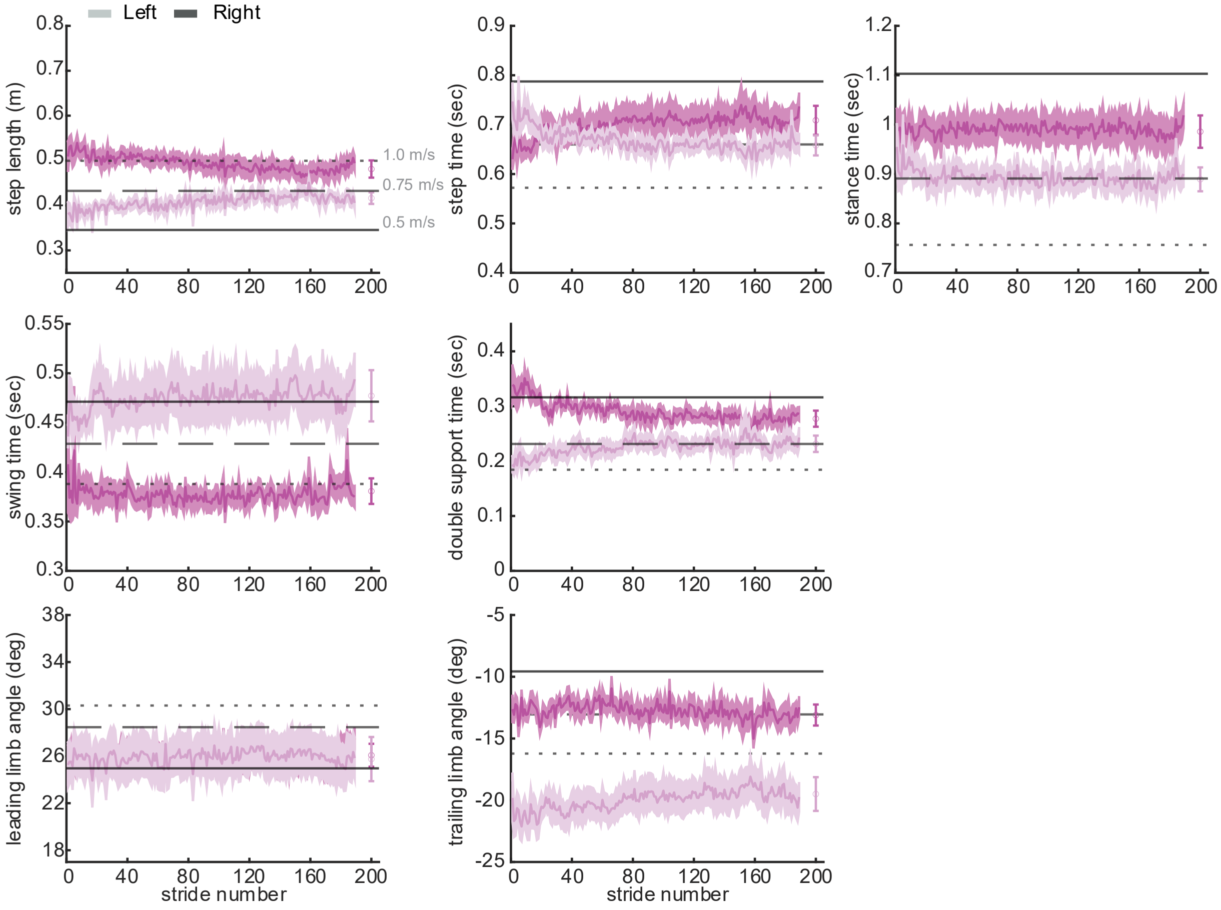


**Figure S2.** Fast spatiotemporal and kinematic traces: Plots depict step-by-step values for step length, step time, stance time, swing time, double support time, leading limb angle, and trailing limb angle within Fast. Lines and shading conventions follow Figure S1. Asymmetry patterns all agree with our previous work[18] (aside for leading limb angle) but are collected at slower treadmill speeds. Baseline walking values at each speed are shown by horizontal lines (1.0 m/s: dotted; 0.75 m/s: dashed; 0.5 m/s: solid).

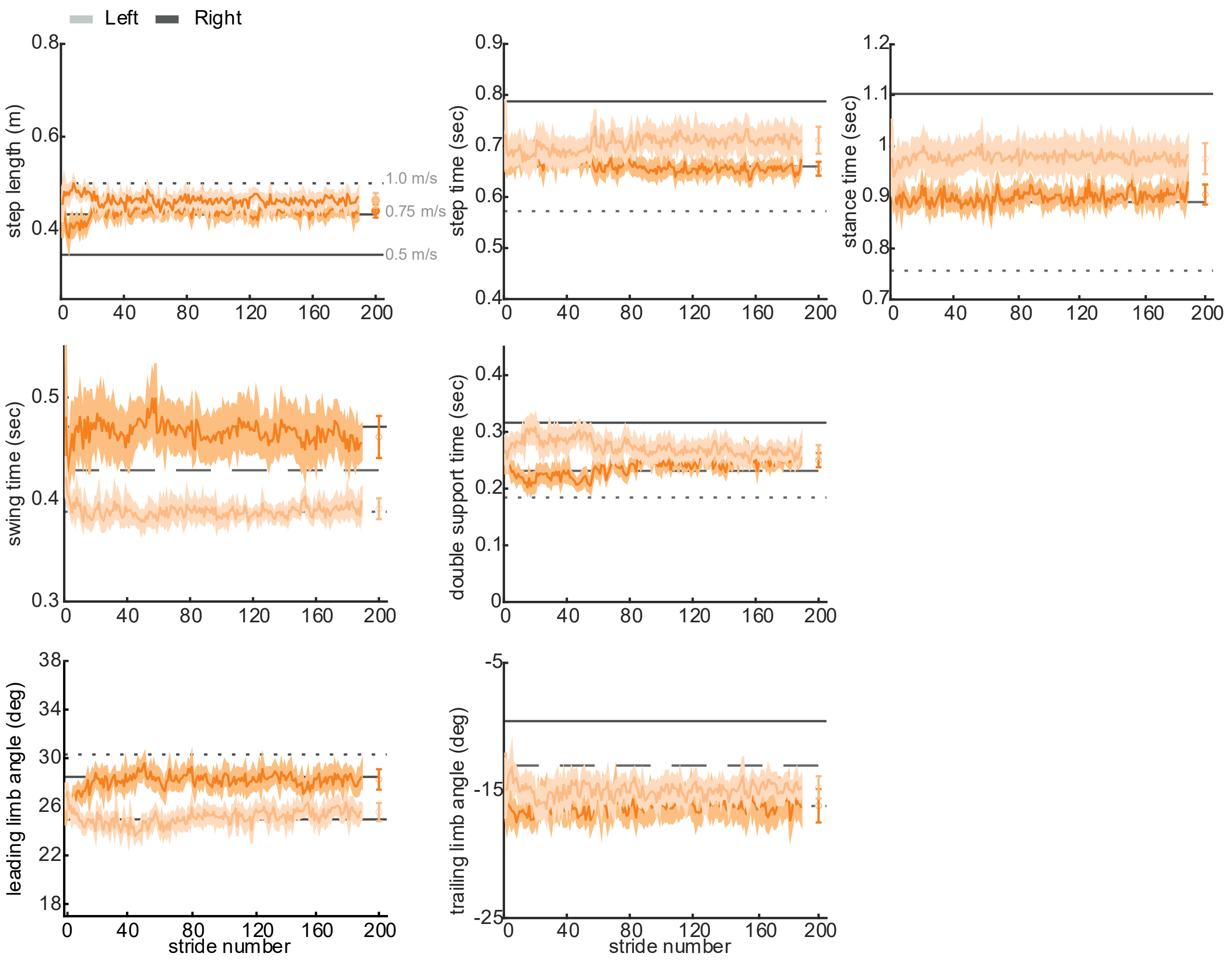


**Figure S3.** Accelerate spatiotemporal and kinematic traces: Plots depict step-by-step values for step length, step time, stance time, swing time, double support time, leading limb angle, and trailing limb angle during Accelerate. **Lines and shading conventions follow Figure S1.** Asymmetry patterns all agree with our previous work[18] but are collected at slower treadmill speeds. Baseline walking values at each speed are shown by horizontal lines (1.0 m/s: dotted; 0.75 m/s: dashed; 0.5 m/s: solid).

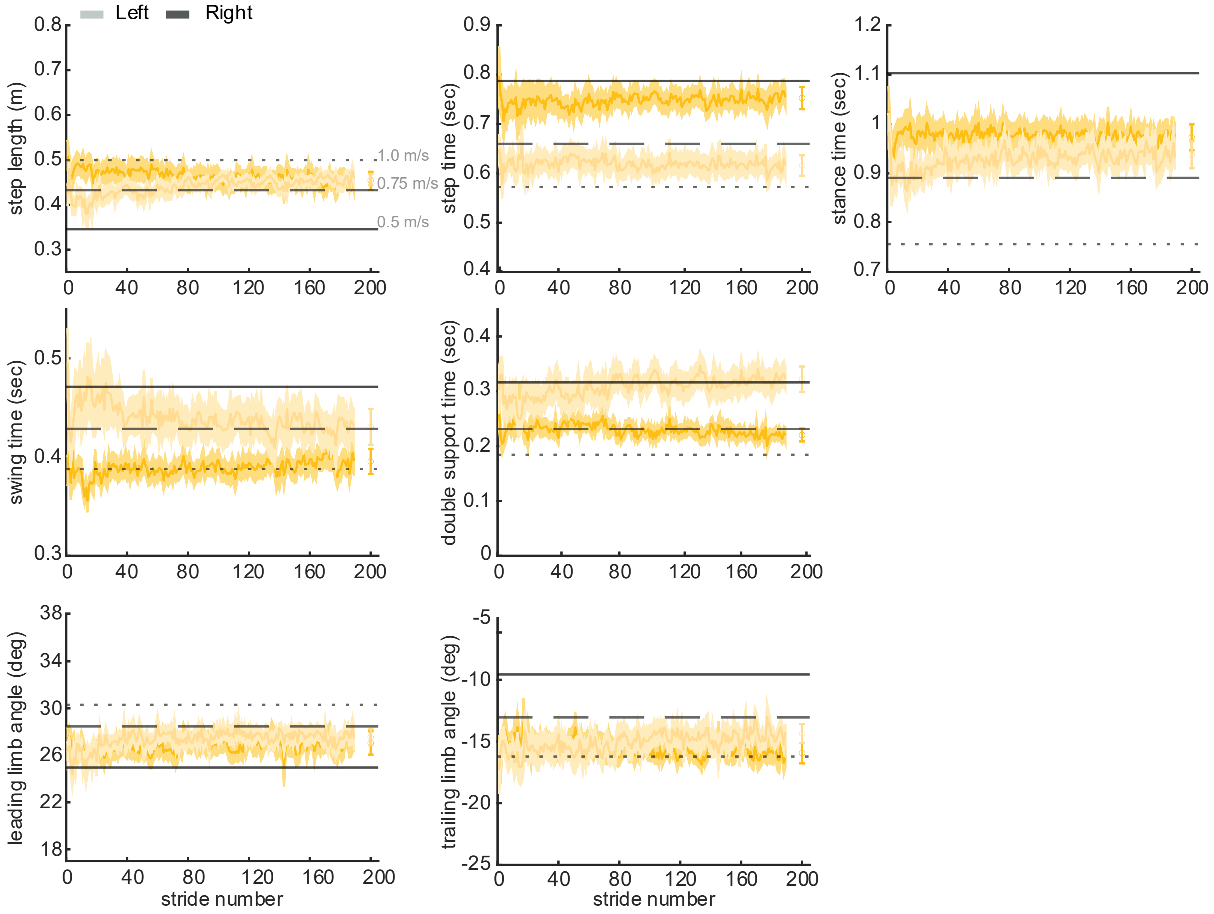


**Figure S4.** Decelerate spatiotemporal and kinematic traces: Plots depict step-by-step values for step length, step time, stance time, swing time, double support time, leading limb angle, and trailing limb angle in Decelerate. Lines and shading conventions are consistent with Figure S1. Asymmetry patterns all agree with our previous work[18] but are collected at slower treadmill speeds. Baseline walking values at each speed are shown by horizontal lines (1.0 m/s: dotted; 0.75 m/s: dashed; 0.5 m/s: solid).
